# Measurement uncertainty matters: ecological management using POMDPs

**DOI:** 10.1101/055319

**Authors:** Milad Memarzadeh, Carl Boettiger

**Affiliations:** ESPM Dept, University of California Berkeley, Berkeley CA, USA

## Abstract

Over the past 30 years, researchers have used various approximations to address the impact of measurement uncertainty on optimal management policy.This literature has consistently suggested the counter-intuitive proposition that increasing harvest levels in the presence of measurement error is often optimal. Here, we use state-of-the art algorithms for Partially Observed Markov Decision Processes (POMDPs) to provide the first complete solution to this classic problem, and demonstrate that contrary to previous work, the resulting policy is usually more conservative than without measurement error. We demonstrate that management which underestimates the measurement error results in both low economic returns and high frequency of stock collapses, while overestimating the role of measurement error can still result in nearly-optimal economic performance while avoiding stock collapse.

## Introduction

Measuring and understanding collapse and over-exploitation of fisheries world-wide remains an essential issue for the ecological sciences (Costello *et al.* 2016). Effective management of marine ecosystems also often informs conservation and resource management in less well-studied systems. The principle of constant escapement-that the optimal management policy of a renewable resource such as a fish stock is accomplished by harvesting the stock down to a fixed population size-underlies much of the research and management in fisheries, dating back to early work by Beverton & Holt (1957) and Schaefer (1957) which proved this principle in deterministic models. In 1979, Reed (1979) extended this work with a classic proof demonstrating that this principle still holds even in stochastic models where the future growth of a population involves a random element. In Reed’s construction, the manager measures the stock size without error each time step before setting the harvest policy. An important consequence of this construction is that this model can thus never lead to accidental extinction of the stock.

Researchers have long recognized the weakness of this assumption of perfect knowledge of the stock size and have sought to understand how it may impact the resulting management policy. Because there is no risk of accidental extinction in this scenario, it is possible that the optimal policy may be more aggressive than is warranted. Unfortunately, due to the complexity of the analysis required to account for measurement uncertainty, this question has never been satisfactorily answered. Moreover, previous attempts at an approximate solution have consistently found the surprising conclusion that the Reed solution is usually less aggressive (dictating higher escapement or lower catch quotas) than attempts to account for measurement uncertainty. We briefly review the main results in the ecological and resource economics literature that have attempted to address this through various approximations before presenting our solution, which draws on substantial algorithmic advances that have been made in other fields.

The important way in which perfect observations diminish the role of uncertainty in the model of Reed (1979) was first recognized in the classic work of Clark & Kirkwood (1986). Clark & Kirkwood (1986) acknowledge the importance of measurement uncertainty would have on the optimal solution, while observing that if this assumption were relaxed, the “difficulty of the problem increases markedly.” Thus, rather than address measurement error question head-on, Clark & Kirkwood (1986) chooses a clever and more nuanced approach to increase the impact that the inherently stochastic growth process of the model has on the optimal solution. In the Reed formulation, harvest occurs after the stock is measured and before the stochastic recruitment takes place. As a result, the population cannot go extinct, unless by design, as the manager can guarantee the choice of harvest is less than the total stock size. Clark & Kirkwood (1986) study a slight modification of this model, in which stock size is still measured without error, but the stochastic recruitment process occurs before harvest takes place. Under this scenario, extinction is possible if realized growth falls sufficiently short of expected growth that harvest exceeds the stock size. In this scenario, Clark & Kirkwood (1986) demonstrate that the optimal solution deviates from the constant escapement rule. Surprisingly, the resulting optimal policy is sometimes less cautious than the constant-escapement policy. In certain cases, Clark & Kirkwood (1986) demonstrate that harvesting to extinction is actually optimal.

Roughgarden & Smith (1996) take a complementary approach, introducing measurement uncertainty but searching only for the optimal constant-escapement, despite presenting any evidence (and indeed contrary to Clark & Kirkwood (1986)) that the optimal solution under this circumstance should be a constant-escapement policy. Roughgarden & Smith (1996) also extend the complexity one step further by also introducing the potential for implementation uncertainty-that the realized harvest can differ from the proposed catch quota by some random shock. By applying the rule-of-thumb assumption that the optimal policy can be expressed simply as a constant escapement (or equivalently, target-stock), the set of possible policies is reduced dramatically and the calculation is relatively trivial. As expected, this restriction results in a policy that is always more conservative than the Reed solution which ignores measurement error, but both economically inefficient and, as we shall see later, still less conservative than the truly optimal policy for large stocks. Sethi *et al.* (2005) seek to address the limitations of Roughgarden & Smith (1996) by searching for the true optimal policy without the strong assumption of constant escapement.

Like Clark & Kirkwood (1986), Sethi *et al.* (2005) finds support for the counter-intuitive result of an optimal policy that is significantly less cautious when accounting for measurement uncertainty than the corresponding constant-escapement solution which ignores it. While noting it is difficult to provide a precise analytic justification for this, the authors offer the intuition that the presence of measurement uncertainty results in a positive probability of the stock exceeding the escapement threshold, relative to the same observation made with perfect knowledge that the the stock is below the constant escapement.

Unfortunately, the approach taken by Sethi *et al.* (2005) once again avoids solving the full optimal decisionmaking problem by means of a simplification which we might term the amnesic assumption: during each subsequent decision the manager forgets any previous knowledge of the system. This allows them to use the very same Stochastic Dynamic Programming (SDP) solution method used by Reed (1979) and Clark & Kirkwood (1986). This method assumes that problem in question is a Markov Decision Process (MDP): one in which given the state *x* and action *a,* the probability of a transition to state *x′;* is conditionally independent of all previous states and actions (the Markov property). This works only if the state is known without error-once we introduce any amount of measurement uncertainty about the state *x*, the process with respect to the observed state is no longer Markovian but depends on all previous observations. As we shall discuss in detail later, this problem is solved by considering the ‘belief’ state rather than the observed state, which restores the Markov property (at the cost of much greater problem complexity). Sethi *et al.* (2005) ignore this and proceed as follows:

The transition matrix between observed state in the present to the observed state in the next time interval is determined by integrating over the probability that the true stock size is *x* given a measurement *m*, but this probability is unknown. Their model formulation specifies only the probability of observing a value *m* given the true stock is *x*; not the inverse. That inverse is given by Bayes Law, which depends on knowledge of a prior on the true stock size, *P*(*x*). Technically such a prior should reflect the manager’s prior knowledge of the stock, which would violate the Markovian assumption and preclude an SDP solution. Sethi *et al.* (2005) sidestep this by implicitly assuming this prior is fixed independent of previous observations-effectively forgetting the history of previous stock sizes. This assumption of amnesia restores the Markov property and permits the SDP solution. It is also this assumption that is thus responsible for their counter-intuitive result that accounting for measurement uncertainty should lead to a more aggressive harvest.

Here we present a solution to this problem that addresses the complexity first recognized by Clark & Kirkwood (1986) head on, avoiding the simplifying assumptions required by Roughgarden & Smith (1996) and Sethi *et al.* (2005). To do so, we build upon methods first developed in the artificial intelligence literature about the same time as Reed’s original work where much progress has been made recently in the discovery of computationally efficient algorithms practical even for large problems. This approach is known as methods for solving Partially Observable Markov Decision Processes or POMDPs.

## Model and Methods

### Partially Observable Markov Decision Process

A Partially Observable Markov Decision Process (POMDP) (Smallwood & Sondik 1973; Sondik 1978) isa generalization of a Markov Decision Process (MDP), which is a fundamental model for sequential decision making. In the ecological literature, MDPs are often referred to by their solution method: Stochastic Dynamic Programming (SDP see Marescot *et al.* 2013) The MDP approach has been widely used in behavioral ecology (Mangel & Clark 1988) as well as conservation ecology and resource economics e.g.(Mangel 1985). Introductions to MDP and solving them through dynamic programming are provided in Mangel (1985), Bertsekas (1996) and Sutton & Barto (1998). Specifically, POMDPs overcome one of the main limitations of MDP which is the assumption about the full observability of the system’s state space. In the POMDP framework, the exact state of the system cannot be observed directly, but can be inferred by indirect measurements.

A POMDP is defined by a tuple (*X, A, Z, T, O, R, γ, b_0_,*) where *X* is system’s state, *A* is set of actions available to the agent, *Z* is observations of states, *T* is transition probability function (that defines dynamics of the system), *O* is the emission probability function (that defines precision of measurements), *R* is the reward function, γ is discount factor, and *b_0_* is the initial belief state. System can experience any state *x* among a finite discrete set *X* = {1,2,…, |*X*|}. The agent can select any action *a* among a finite set *A* = {1,2,…, |*A*|} at each time step. Based on the current state and action, she receives reward *R*(*x, a*). Time is discretized in steps, and variables *x_t_*,*a_t_*,*r_t_* indicate state, action and reward at time *t* respectively. Expected reward is assigned by function 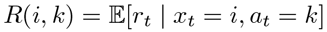,where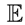 indicates the statistical expectation. After an action is taken, the state evolves stochastically following a Markov process governed by transition probability function 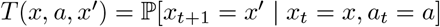, where 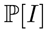 indicates the probability of event *I*. Transition function defines the dynamics of the system.

In MDPs, action *a_t_* follows the full observation of the state *s_t_* that, given the Markovian assumption, is a sufficient statistics for the process, meaning that given the current state, future is independent of the past. On the contrary, POMDPs assume that at each time *t* agent has access only to a noisy and incomplete measure of the current state, *z_t_*, which can assume one value in a finite set *Z* = {1,2,…,|*Z*|}. The precision of the observations is governed by the emission probability function 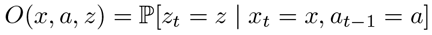. In POMDPs, the agent’s knowledge about the current state is represented by a probability distribution, or belief state, 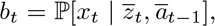, which sets 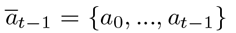 and 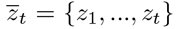 are the history of measurements and actions up to time *t*, respectively. Belief state is a sufficient statistic for all past actions and measurements in POMDP, meaning that given the current belief state, decision making process is independent of the past. Belief state *b_t_* belongs to ***B***, the space of probability distributions over the state of the system ***X***. Belief at time *t* +1 can be updated using the Bayes rule as follow:

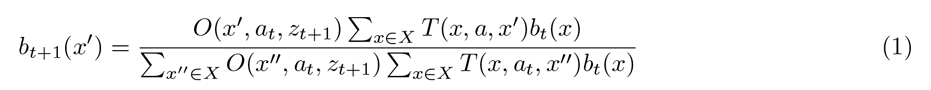

MDPs and POMDPs can be represented as a probabilistic graphical models (Barber (2012)) as in Figure 1 and Figure 2, respectively. circles define random variables, squares decision variables, diamonds utility variables, and arrows the (potential) dependence among variables.

Only shaded variables are observable to the agent. In the context of POMDP, the agent’s goal is to maximize value *V*, defined as the expected sumof discounted rewards over an infinite time horizon: 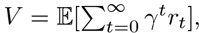using discount factor γ that relates the future rewards to their net present value. The agent’s behavior is defined by a policy, i.e. a mapping between belief state and actions: *π*:*B ⟶ A*. When a policy *π* is adopted, action at time *t* can be selected as *a_t_* = π(*b_t_*). The value depends on policy *π* via the recursive linear equation where *b′* is updated belief *b_t_+_1_,* according to Eq. (1), with *a = a_t_*, *z* = *z_t_+_1_*, b = *b_t_*, and conditional probability 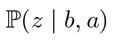 is computedas:

**Figure 1:**
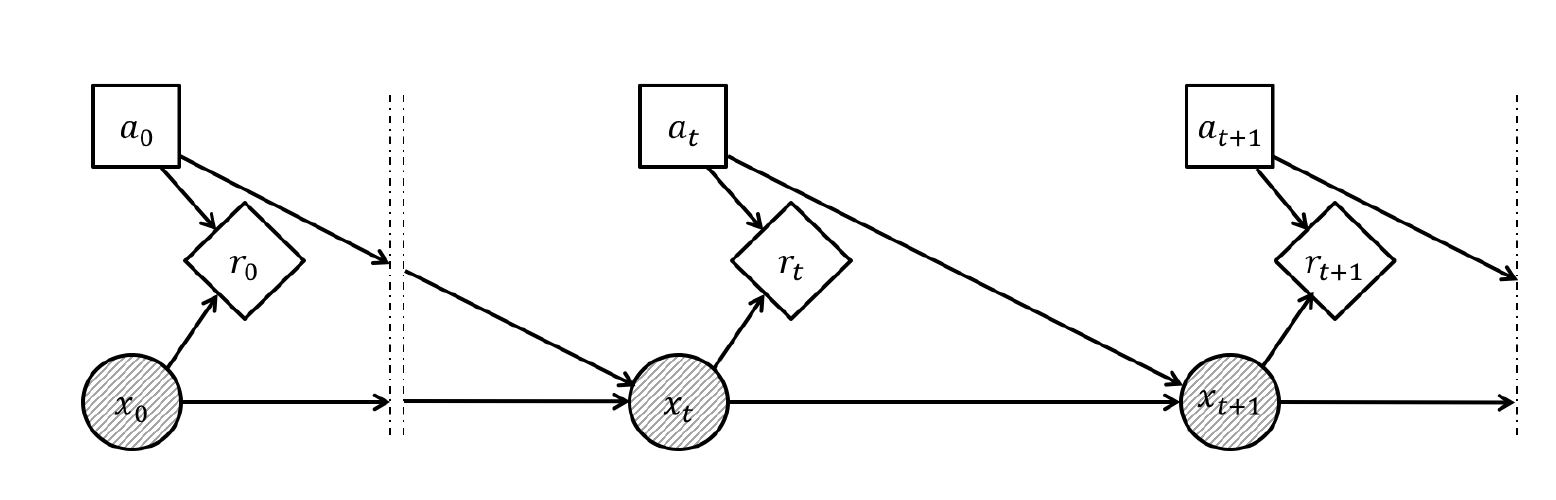
Probabilistic graphical model of MDP

**Figure 2:**
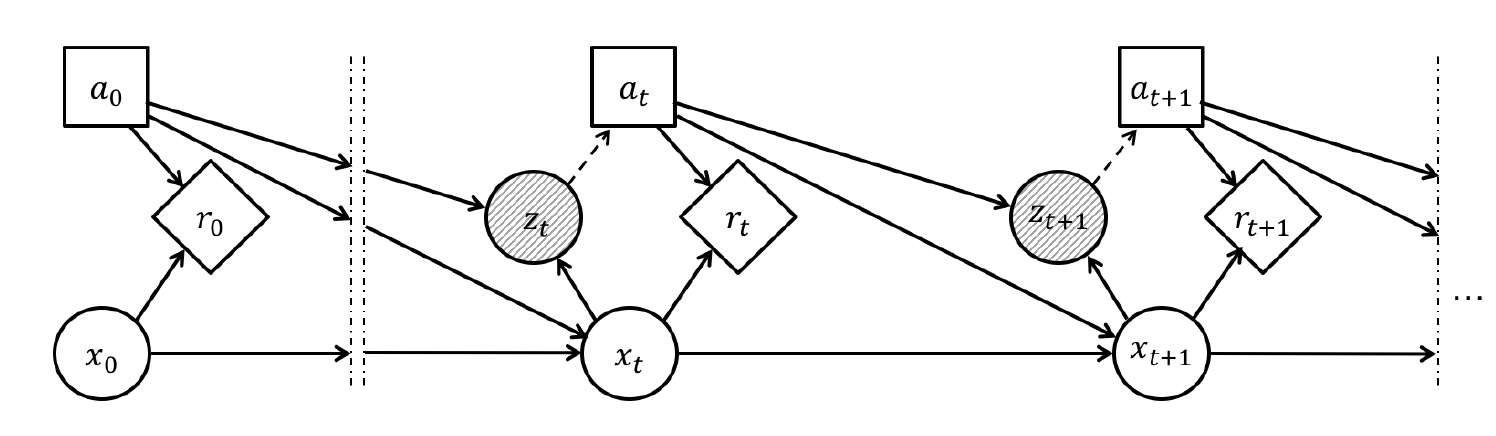
Probabilistic graphical model of POMDP

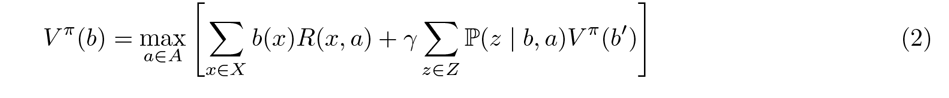

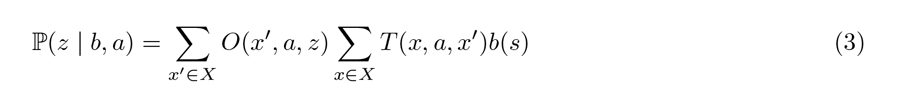

The optimal value is defined by the Bellman equation (Bellman 1957):

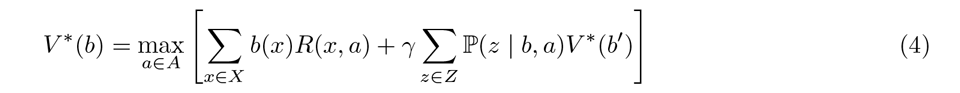

A POMDP problem can be solved by applying the same methods of solving MDP (e.g. value iteration and policy iteration), however as the belief state is defined on an infinite space, the exact solution for the POMDP is generally not available. Kaelbling *et al.* (1998) have proposed witness algorithm for finding exact solution to POMDPs using value iteration, which is only applicable to small systems. Grid-based methods (Lovejoy 1991; Zhou & Hansen 2001) have been successful in the past, however they are not scalable to complex systems due to complexity of the interpolation phase. State-of-the-art methods for solving POMDPs are point-based value iteration methods (Pineau *et al.* 2003) that restrict the search to the beliefs that are reachable from the initial belief state. Generally, the POMDP solution is known to be NP-Hard in the worst case, yet point-based solvers often provide accurate approximations in practice. Throughout this paper, we use Successive Approximations of the Reachable Space under Optimal Policies (SARSOP) (Kurniawati *et al.* 2008), which is an efficient point-based value iteration method and represents state-of-the-art in solving POMDPs both in efficiency and accuracy.

As a supplement to this paper we provide example scripts for replicating the analyses shown here, which can be found at https://github.com/cboettig/appl.

In contrast to MDP, POMDP approaches anywhere in the ecological literature are very few. Previous applications of POMDP to ecological and conservation issues have primarily relied mostly on approaches that can still be solved using traditional SDP techniques by transforming the problem into a belief MDP (Chadès *et al.* 2008, Chadès 2011; Haight & Polasky 2010; Williams 2011; Springborn & Sanchirico 2013). This computationally intensive approach has thus restricted such examples to systems with only a very small number of states and/or actions. By leveraging newer point-based algorithms recently developed in the artificial intelligence community, we are able to adapt these methods to the much larger problem posed by the optimal fisheries harvest problem. We now present the fisheries optimization problem more precisely.

## Model formulation

We examine the performance of proposed method on management of a marine ecosystem. Time is discretized and we consider the problem of infinite horizon, *t* = 0,1, 2*,…,*∞. State of the system, *x_t_* is defined by the fish stock at time *t*. The harvesting policy is formulated as follow:where *a_t_* is the attempted harvest at time step *t*.

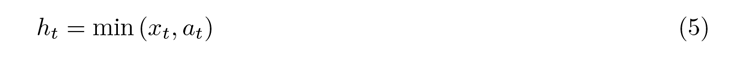

There are two sources of uncertainty: population dynamics (i.e. growth of the stock), and stock measurements.

The population dynamics is formulated as follow:where *x_t_* and *s_t_* are the stock and escapement at time step *t* and *ϵ_g_* is a noise distributed either log-normally or uniformly with mean 1 and standard deviation of *σ_g_*. Eq. (6) corresponds to the transition probability function defined in the POMDP framework: future state of the system, *x_t+1_* is defined as a function of escapement at current time step, *s_t_*, defined as: *s_t_* = *x_t_* - *h_t_*. This analogy is proportional to that of general POMDP framework as the future state of the system, *x_t+1_*, depends on current state, *x_t_*, and current action, *a_t_* (based on Eq. (5)).

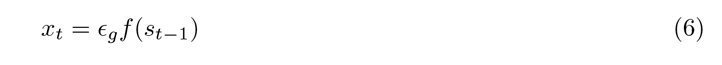

We consider three candidate parametric models for the population dynamics (*f*): the Ricker model (Ricker 1954),the Beverton-Holt model (Beverton & Holt 1957),and the Allen model (Allen *et al.* 2005),where *K* denotes the carrying capacity of the system, *r* the maximum per capita growth rate, and *C* the location of the unstable steady state (i.e. the tipping point).

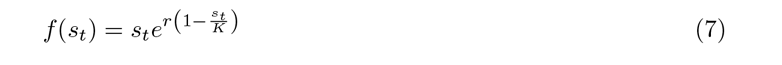

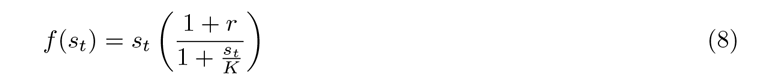

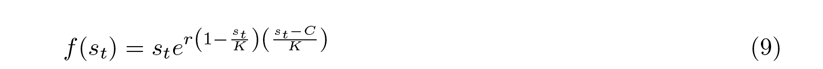

Stock measurement is formulated as follow:where *x_t_* and *z_t_* are the actual stock and stock measurement at time step *t* and *ϵ_m_* is a noise distributed either log-normally or uniformly with mean 1 and standard deviation of σ_m_. Eq. (10) corresponds to the emission probability function defined in the POMDP framework which relates the noisy observations, *z_t_*, to system’s state, *x_t_*. In the general POMDP framework, observations can depend on agent’s actions as well, but in the examples used in this paper, measurements only depend on the states.

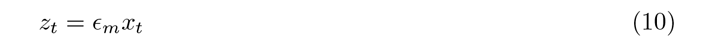

The reward function is formulated as:with 0 ≤ *α* <1. This definition is proportional to the definition of the reward function in POMDP framework, *R*(*x_t_*, *a_t_*), as here the reward at time *t*, *r_t_* depends on the current state, *x_t_*, and current action, *a_t_*, based on the definition of *h_t_* in Eq. (5). Here we assume that reward of harvesting is $1 per harvest, *h_t_* andthe price of harvesting is *α* per action, *a_t_*.

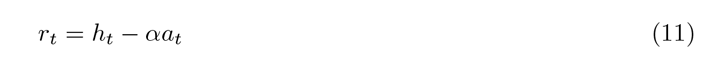

For our focal example, we consider a system of size |*X*| = 50, where the carrying capacity fixed at ***K*** = 33 and the maximum per capita growth rate at *r* =1 for each of the three models. In the Allen model, we fix the location of tipping point is *C =*10. Throughout, we use a discount factor of γ = 0.95. We evaluate the effect of different noise levels in both growth and measurements, *σ_g_*, σ_m_ *ϵ* {0, 0.005, 0.1, 0.5}. In order for uniform and lognormal noises to be comparable (i.e. have the same variance) we define the *µ* and *σ* of lognormal distribution as follows:

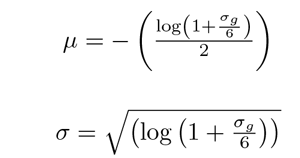

In this manner our *σ* values are directly comparable to those used by Sethi *et al.* (2005), who consider only the uniform noise case. In our supplementary material we consider variations of each of these choices, including the comparable results under the uniform noise distribution.

## Results

To illustrate the impact of measurement error on management outcomes, we compare 500 independent replicate simulations with stochastic growth (*σ_g_*= 0.1) and measurement error present (*σ_m_* = 0.1 and *σ_m_* = 0.9). For each simulation, we compare the outcome from managing under the optimal POMDP solution to that of the POMDP solution which assumes the measurement error is either larger (3a) or smaller (3b) than it really is in the simulation. It can be seen that underestimating the measurement error suggests less conservative harvesting policies, i.e. over-exploiting (Figure 3c) and therefore, will result in dramatic collapse of the system (Figure 3b) and significant loss in revenue (Figure 3a). On the other side, we show that overestimating the measurement error might result in a trivial loss in revenue but does not result in the collapse of the system as the agent takes more conservative actions. In the results presented in Figure 3 the population growth of the system is defined by Ricker model and the noise error is log-normally distributed. The results in the case of uniform noise is presented in the supporting information (Figure A1), which shows a similar pattern. The results in the case of Beverton-Holt model is also the same and is reported in the supporting information (Figure A2). We also show that defining the measurement error as log-normal is significantly better than uniform (Figure A3 and A4). The log-normal noise has a larger tail and represents more realistic noise, and hence will result on adaptation of more conservative harvesting policy.

**Figure 3:**
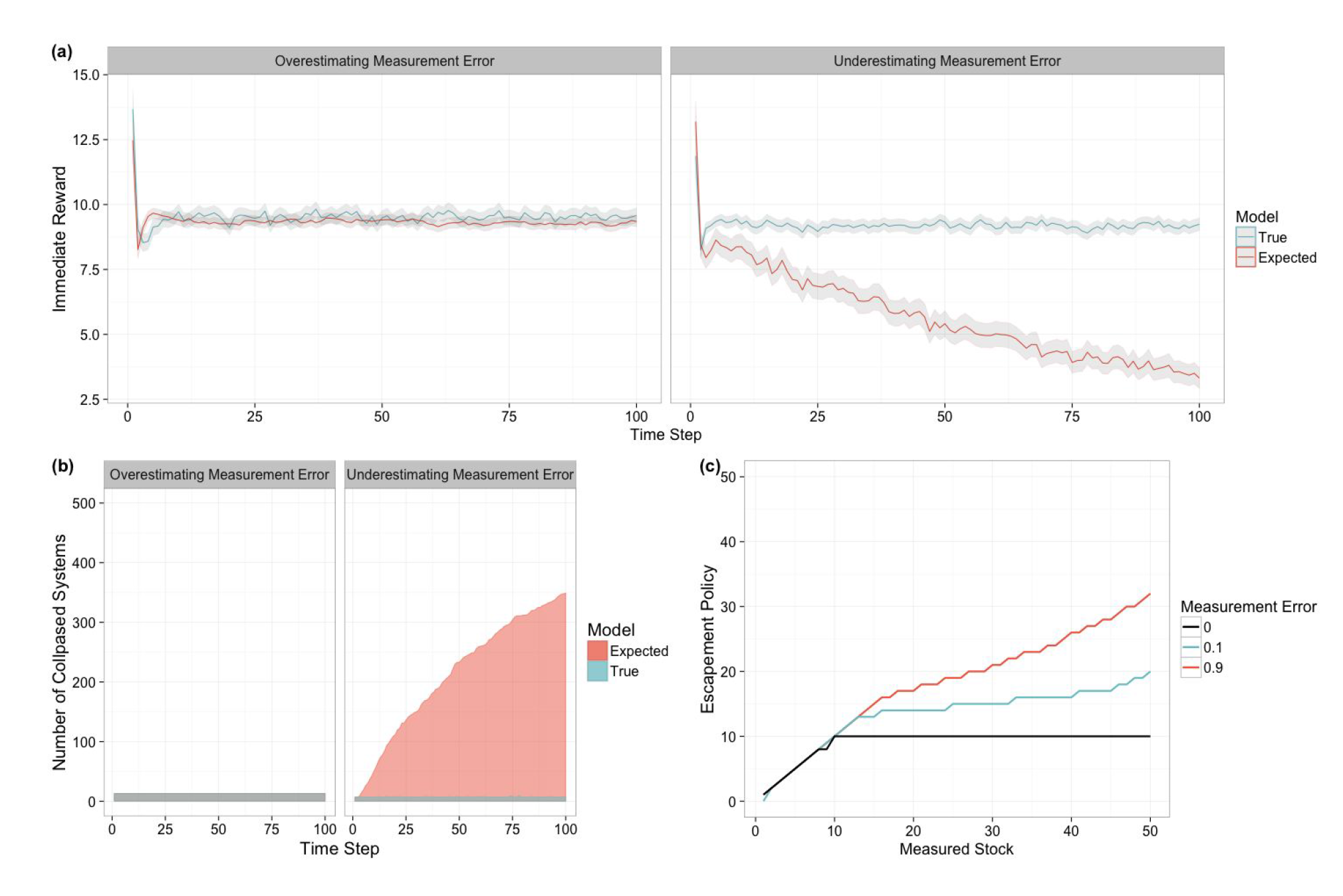
Results of 500 independent forward simulations of managing a marine ecosystem comparing overestimating measurement error with underestimating it: (a) immediate reward that agent receives through the management process, (b) number of collapsed systems in time, and (c) optimal escapement policies for different measurement errors. The growth model is Ricker model in this figure, the noise is lognormally distributed, the carrying capacity of the system is *K =*33, the maximum per capita growth rate is *r* = 1, and the discount factor is *γ* = 0.95. Figure 4 shows the optimal escapement policy under different growth models, growth error, and measurement error. The vertical dashed line represents the carrying capacity of the system and all noises are distributed log-normally. Once measurement noise is introduced, the resulting optimal escapement policy is no longer the traditionally-known fixed piecewise-linear policy. The growth error has a significant effect only in the case of Allen model, while in other cases, the effect of growth error is trivial. Although, the measurement error has significant effect on the policy. As the measurement noise increases, the policy becomes more and more conservative with the exception in the case of Allen model with large measurement error. Existence of the tipping point (located at *x* = 10) makes the adapted policy more conservative below the tipping point, less conservative after tipping point, and again more conservative in large stock sizes.

**Figure 4:**
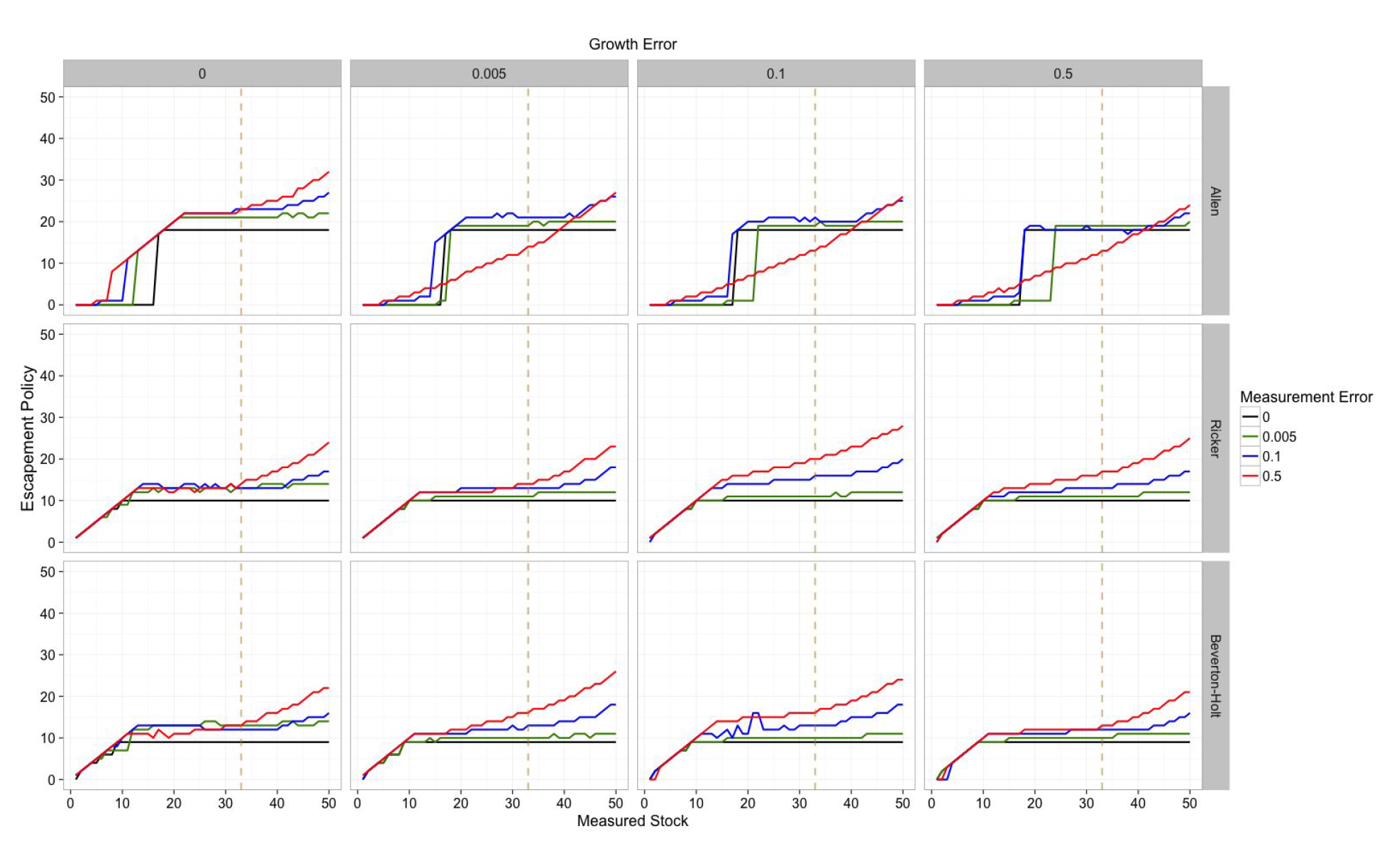
Optimal escapement policy for different growth models, growth error, and measurement errors (log-normal noise). The dashed vertical line represents the carrying capacity of the system, *K* = 33. State space is discretized into |*S*| = 50 states, the maximum per capita growth rate is *r* = 1, and the discount factor is *γ* = 0.95.

We evaluated the effect of modified reward function (with addition of harvesting cost, *α* > 0 in (Eq. 11) to the adapted escapement policy (shown in Figure A6 in the supporting information). Agent adapts a slightly more conservative policy, but we find the effect to be trivial. Figure A7 shows a similar pattern with uniform noise.

We simulate 1000 forward runs of controlling the above-mentioned marine ecosystem and show the density of visited states through time in Figure 5. It is worth noticing that the density of collapse state is not shown in the figure. The vertical dashed line represents the stable state that the agent preserves the system in for the case of no uncertainty in growth. All models tend to preserve the system in a state below the carrying capacity (*s* = 20 in cases of Allen and Ricker models,and *s* = 14 in the case of Beverton-Holt model). Introduction of the measurement error, shifts the stable state to larger stocks, as the agent tries to keep the system in safer state, because presence of high error in measurements might result in collapse of the system. We show the case of uniform error in the supporting information (Figure A8), that agent keeps the system in larger stocks compared to the case of no growth noise, which is again as a result of assigning high probability to the states far from the true state by uniform noise.

**Figure 5:**
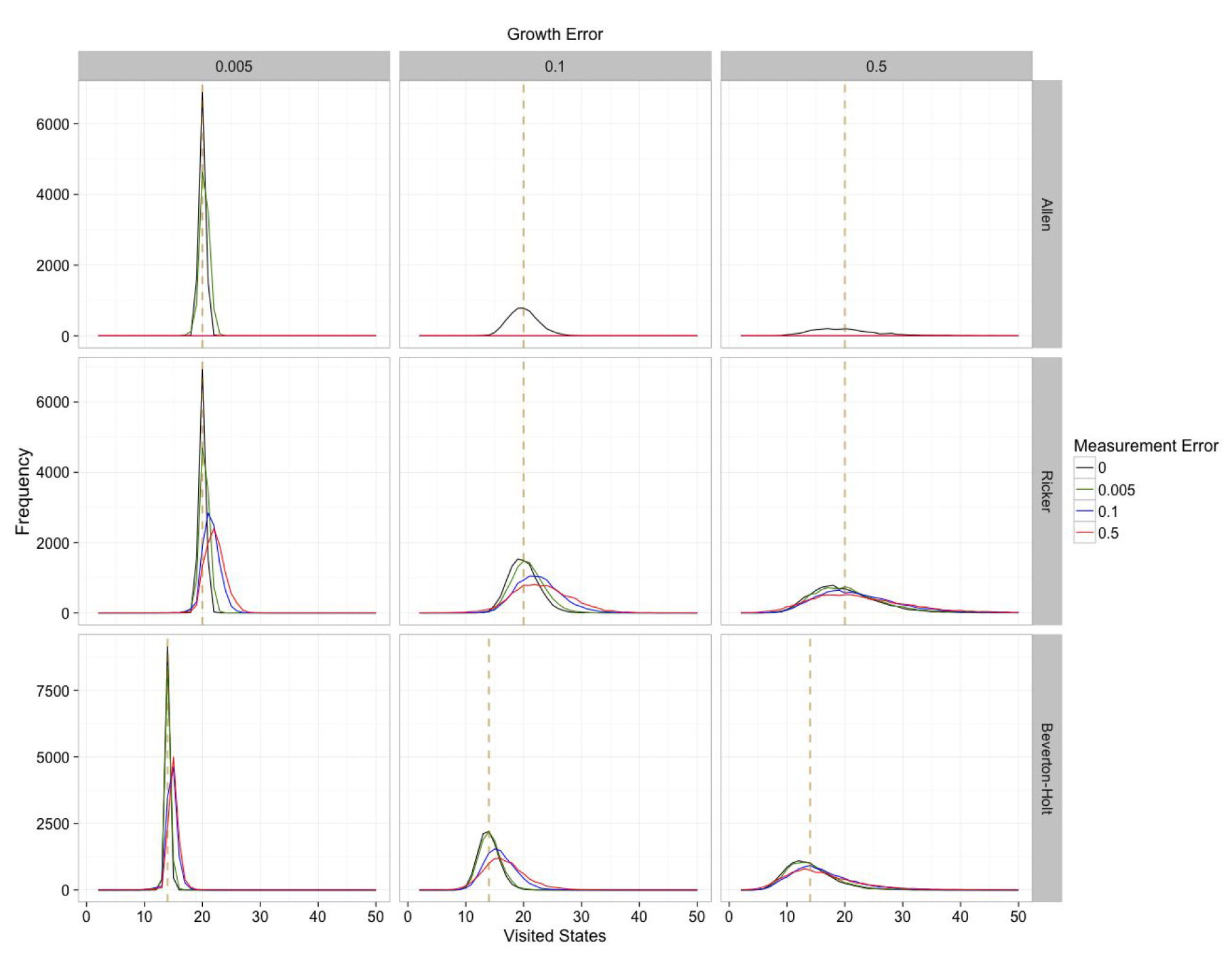
Density of visited states in 1000 independent simulations of managing the mentioned marine ecosystem for different growth models, growth error, and measurement errors (log-normal noise). The dashed vertical line represents the stable state that agent keeps the system in for the case of no uncertainty

Unlike stochastic growth which is an intrinsic part of the ecological system (arising from random demographic events of birth and death or the natural fluctuations of environment), measurement uncertainty can potentially be reduced directly by acts of the manager. Consequently, it is of interest to understand the economic value that could be derived from a given reduction in the measurement error. Figure 6 shows the relative difference in value of the provided measurements with respect to perfect measurements, which we generally call relative Value of Information (VoI). It is defined as 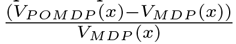, where *V_POMDP_*(*x*) is the value of the POMDP problem with the corresponding measurement and growth error and *V_MDP_*(*x*) is the value of MDP problem with the corresponding growth error.The dashed vertical line represents the carrying capacity of the system.

## Discussion

We have shown that fully accounting for measurement error in calculating the optimal fishing harvest frequently leads to a policy that is more conservative (harvesting less) as the measurement error increases. The importance of this intuitively sensible conclusion is underscored by the contrast to both much of current fisheries management practice and previous research into measurement uncertainty. Most fishing policy still reflects the constant-escapement principle that Reed (1979) derives in the absence of measurement error, which we have shown leads to over-harvesting. Meanwhile, previous work on the role of measurement uncertainty (Clark & Kirkwood 1986; Sethi *et al.* 2005) has relied on approximations which have argued that accounting for measurement uncertainty should result in an *increased* harvest (lower target stock size). The danger, both economic and environmental, of this assumption can clearly be seen in the comparisons in Figure 3. Underestimating measurement uncertainty (3b) results in an economic net value that is only 1/3rd of the optimal, and widespread collapse of the stock, with nearly 70% of simulations collapsing to extinction by the 100th time step in our example. A policy which ignored uncertainty all-together or worse, proposed an even higher harvest, would have even worse results. Meanwhile, overestimating the measurement uncertainty (3a) not only avoids stock collapse but results in an economic value from the stock that is consistently near the optimal value.

**Figure 6:**
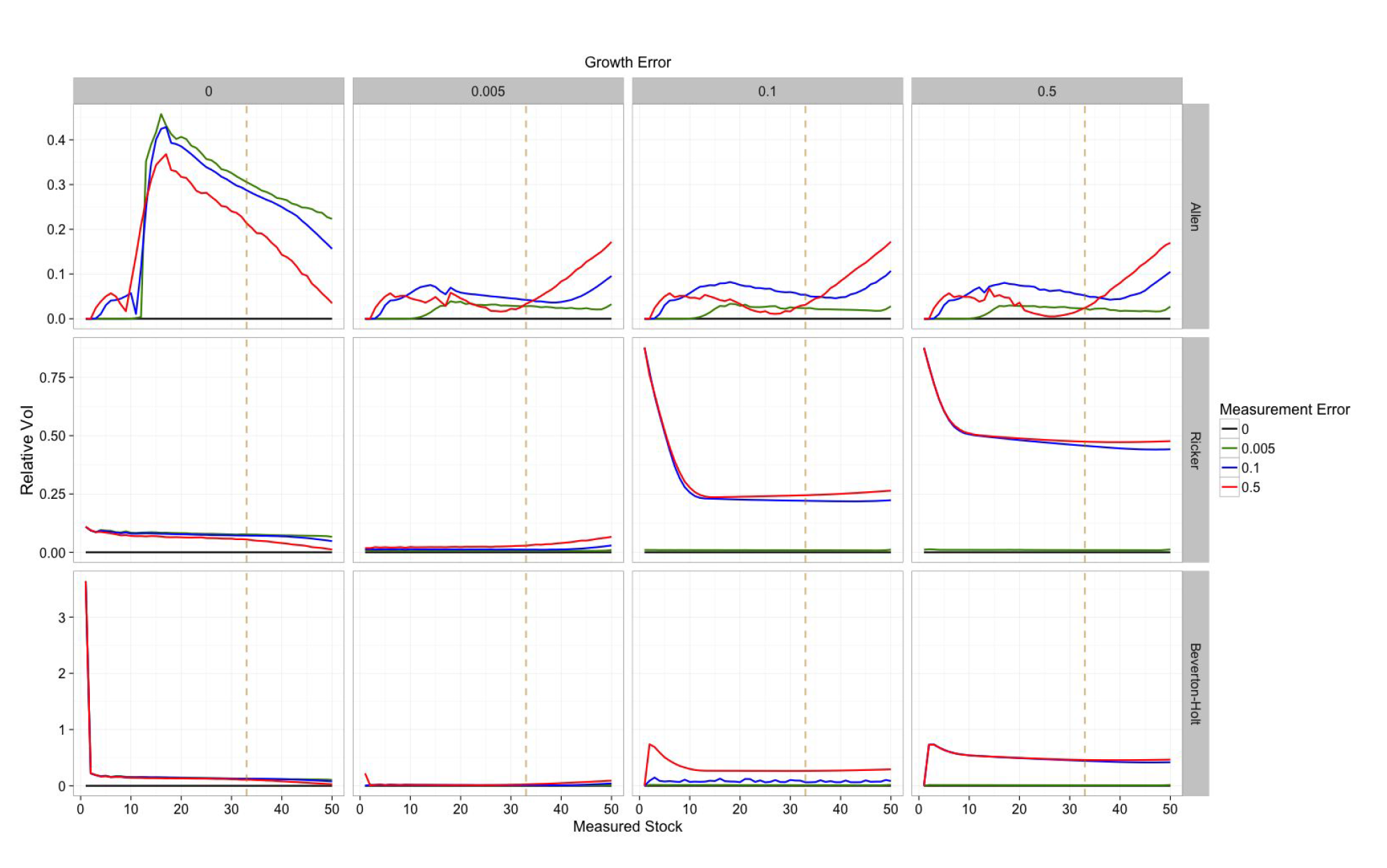
Relative value of information for different growth models, growth error, and measurement errors (log-normal noise)

Note that in this model we have assigned no intrinsic value to preserving the stock, apart from it’s future economic potential for harvest. Additionally, we have made the simplifying assumption that there is no cost to fishing and no impact on the price per unit of fish from a large harvest compared to a small one. While we consider relaxing these assumptions in the appendix in favor of a more realistic reward function, it is important to note that each of these amounts to a ‘worst-case’ scenario:all result in a potentially more aggressive fishing policy than would if they were relaxed, since either a cost to additional fishing effort or decrease in price would provide reason for a smaller harvest. Consequently, the more precautionary policies we find here are not driven by economic assumptions.

The ability of our approach to capture nearly optimal economic returns even while overestimating the total measurement uncertainty reflects the value of nuanced optimal policy accounting for measurement error explicitly rather than the a trivially precautionary approach such as proposed in Roughgarden & Smith (1996). The rule-of-thumb calculated in Roughgarden & Smith (1996) argues that optimal escapement should always be set at 3/4 of the unharvested equilibrium stock size, in part to reflect the risk of measurement error. Our results (e.g. Figure 4) suggest an optimum solution that is sometimes less conservative than this, while still more conservative than the solution ignoring measurement error. For certain stock values our solution is in fact more conservative than this rule of thumb, particularly in the case of very large measured stock values. Stock values above the actual carrying capacity are exceedingly unlikely (Figure 5), and thus these large estimates are particularly likely to be the chance results of high measurement uncertainty and treated cautiously. Hence the strong rise in constant escapement at the very high-stock end of our policy curves (Figure 3). This underscores the importance of having an optimal policy that depends explicitly on the estimated stock size rather than a constant target stock escapement.

Our result is not always strictly more conservative. Under the less biologically realistic assumption of uniform noise, we find a policy that is less conservative at low stock sizes (Figure A7), though still more conservative at high stock sizes. This is similar to the results of Sethi *et al.* (2005) who also examine the uniform noise case, though even for comparable levels of measurement and growth uncertainty in the uniform model, our exact solution still predicts a higher escapement than their approximation provides. In the special case of the Allen model where the stock dynamics contain an explicit tipping point, we find that it is sometimes optimal to implement a policy that is less conservative than had measurement error been ignored. This reflects the extreme difficulty in managing a system with tipping points under a high degree of stochastic growth and measurement error. Even under optimal management or the harvest-free scenario, stocks under this model have a significant chance of collapsing. In face of this risk, our profit-driven optimization frequently decides that it may be better to harvest the entire stock while it still exists than to risk it going extinct on its own. Still, under suitably small noise where random extinction is less likely, we find harvest policies proposed that are more intermediate, sometimes more conservative and sometimes less conservative.

This difference between the Allen model and the other models is also reflected in the value of information (Figure 6), which is highest at low stock sizes for the models without a tipping point such as Ricker and Beverton-Holt, but highest in the vicinity the tipping point for the Allen model. Other than this difference for the Allen tipping point model, we have shown that our result is generally robust to our assumptions: this pattern holds for differing models (Ricker, Beverton-Holt), differing magnitudes and also different shapes of noise distributions (log-normal, as considered by Clark & Kirkwood (1986) or uniform, considered by Sethi *et al.* (2005)), the choice of the reward/profit equation, or the details of the discretization. This robustness is not surprising, being consistent with sensitivity observed in previous work. Indeed, in the absence of measurement error most of these scenarios were already covered by Reed’s proof, with the exception of the Allen model whose change in concavity violates the theorem. Thus it is perhaps all the more surprising that such results are not robust to the assumption of measurement error.

This important qualitative impact that measurement error has on the optimal solution of an important and well-studied ecological management problem has more general implications. The role of measurement error is most often acknowledged in the breach - often a mathematically necessary assumption one hopes will have little qualitative impact, at least so long as those measurement errors are unbiased and the data is not too sparse. After all, inherent stochasticity has often been found to make only small differences from deterministic solutions, as Reed (1979) proved in this case. Our result demonstrates how frail such an assumption regarding measurement uncertainty may be, particularly when determining optimal policy. No doubt the mathematical and computational difficulty of accounting for measurement uncertainty has forestalled greater attention to this issue. The growing wave of available data makes such concerns about measurement error all the more pressing, as imperfect but readily available observations make such data-driven optimization possible in more applications. We hope that the recent advances in efficient and accurate POMDP solvers, together with greater attention to these challenges, will encourage a wider consideration of the role of measurement error in determining appropriately precautionary management policy.

In conclusion, we have demonstrated the importance of measurement error in the long-standing question of optimal harvests in fisheries. We have shown that over-estimating the measurement error can still result in policies that are nearly optimal both economically and ecologically, while ignoring or underestimating the measurement noise can cause dramatic collapses of fish stocks even when no tipping point is present. This suggests that the failure to account for measurement error in this way in even the most advanced fisheries management could contribute to long-term trends in declining stocks (e.g. Costello *et al.* 2016). We believe the approach taken here both highlights the importance of accounting for measurement error and provides an example of how this can be accomplished in the POMDP framework using modern algorithms even on complex problems.

## Acknolwdegements

CB acknowledges helpful discussions on this topic with Iadine Chades, Jim Sanchirico, Mike Springsborn, and Perry de Valpine; funding from the UC Berkeley ESPM dept, and computational resources from XSEDE Jetstream [Stewart *et al.* (2015); award DEB160003] and Chameleon testbed supported by the National Science Foundation, Awards 1419141 and 1419152.

